# Quantifying point-mutations in shotgun metagenomic data

**DOI:** 10.1101/438572

**Authors:** Shruthi Magesh, Viktor Jonsson, Johan Bengtsson-Palme

## Abstract

Metagenomics has emerged as a central technique for studying the structure and function of microbial communities. Often the functional analysis is restricted to classification into broad functional categories. However, important phenotypic differences, such as resistance to antibiotics, are often the result of just one or a few point mutations in otherwise identical sequences. Bioinformatic methods for metagenomic analysis have generally been poor at accounting for this fact, resulting in a somewhat limited picture of important aspects of microbial communities. Here, we address this problem by providing a software tool called Mumame, which can distinguish between wildtype and mutated sequences in shotgun metagenomic data and quantify their relative abundances. We demonstrate the utility of the tool by quantifying antibiotic resistance mutations in several publicly available metagenomic data sets. We also identified that sequencing depth is a key factor to detect rare mutations. Therefore, much larger numbers of sequences may be required for reliable detection of mutations than for most other applications of shotgun metagenomics. Mumame is freely available from http://microbiology.se/software/mumame

## Introduction

The revolution in sequencing capacity has created an unprecedented ability to glimpse into the functionality of microbial communities, using large-scale shotgun metagenomic techniques (Quince et al. 2017). This has yielded important insights into broad functional patterns of microbial consortia (Yooseph et al. 2007; Human Microbiome Project Consortium 2012; Sunagawa et al. 2015). However, while overall pathway abundances inferred from metagenomic data can tell us much about the general functions of communities and how they change with e.g. environmental gradients (Bengtsson-Palme 2018; Bahram et al. 2018), there are many important functional differences that are hidden in the subtleties of these communities (Österlund et al. 2017). For example, many antibiotic resistance phenotypes are the results of single point mutations rather than acquisition of novel pathways or genes (Johnning et al. 2013). This complicates the studies of selection pressures in environmental communities, as analysis of such mutations is generally limited to a narrow range of species (Johnning et al. 2015b; Johnning et al. 2015a; Kraupner et al. 2018).

Because of the immense increase in available sequence data, it would be desirable to study these mutations from shotgun metagenomic libraries, much as other traits have been studied at a large scale (Pal et al. 2016). However, attempts to quantify point mutations in metagenomic sequencing data often go wrong because the methods do not sufficiently well distinguish between mutated and wildtype variants of the same gene. For example, a sequenced read may map to a region identical in the mutated and wildtype variant of a gene, causing problems for quantifying their relative proportions (Bengtsson-Palme et al. 2017). In addition, because the sought-after mutations generally are rare in most types of sample, and metagenomic studies are often under-sampled in terms of replicates (Jonsson et al. 2016a), commonly applied statistics methods may not be sufficiently sensitive to reliably detect differences between samples (Jonsson et al. 2016b).

In this study, we attempt to provide a partial remedy to these problems through the introduction of a software tool – Mumame – that can quantify and distinguish between wildtype and mutated gene variants in metagenomic data, and through suggesting a statistical framework for handling the output data of the software. We further demonstrate the ability of the method to detect relevant differences between environmental sample types, estimate the sequencing depths required for the method to perform reliably through simulations, and exemplify the utility of the software on detecting resistance mutations in publicly available metagenomes. The Mumame software package is open-source and freely available from http://microbiology.se/software/mumame

## Methods

### Software implementation

Mumame is implemented in Perl and consists of two commands: mumame, which performs mapping to database of mutations, and mumame_build with builds the database for the former command. The mumame_build command takes a FASTA sequence file and a list of mutations (CSV format) as input. For each entry in the mutation list, it finds the corresponding sequence(s) in the FASTA file, either by sequence identifier or by CARD ARO accessions (Jia et al. 2016). It then excerpts a number of residues upstream and downstream of the mutation position (by default 20 residues for proteins and 55 for nucleotide sequences) and creates one wildtype version and one mutated version of the sequence excerpt with unique sequence IDs. For cases where multiple mutations can occur close to each other on the same sequence, the software attempts to create all possible combinations of mutations (if memory permits – is some situations this is not possible because the number of combinations increase exponentially). The software tool also generates a mapping file between sequence IDs in the database and mutation information from the list.

The main mumame command takes any number of input files containing DNA sequence reads in FASTA or FASTQ format and maps those against the Mumame database using Usearch (Edgar 2010). For this mapping, the software runs Usearch in search_global mode with target coverage set to 0.55 (by default; any value ≥ 0.51 should be feasible for target coverage). The output is then mapped to the wildtype or mutation information in the Mumame database, and data is collected for each input file and combined into one single output table.

The output table generated by Mumame can then be analyzed using the R script (R Core Team 2016) supplied with the Mumame package. The script reads the read counts for all mutation positions detected, both for wildtype and mutated sequences, and assesses if there are significantly different proportions of mutations between different sample groups directly through a generalized linear model. Alternatively, an overdispersed Poisson generalized linear model accounting for the discrete nature of the data and the differences in sequencing depth can be used (Jonsson et al. 2016b; Bengtsson-Palme et al. 2017). The Poisson model is preferable when the number of counts for a targeted gene is low in all sample groups.

### Quantification of mutations in metagenomes

To quantify the abundances of fluoroquinolone resistance mutations in the *gyrA* and *parC* genes (Johnning et al. 2015b), we downloaded the CARD database on 2018-05-24 (Jia et al. 2016). We extracted all mutation information regarding the *gyrA* and *parC* genes from the “snps.txt” file and created a new file with that information. We then created a new Mumame database, with the following command: “mumame_build −i card-data/protein_fasta_protein_variant_model.fasta −m gyrA_parC_snps.txt −o gyrA_parC”. That database was used to map all the reads from the samples generated by Kraupner et al. (2018) to the database using Mumame in the Usearch mode (Edgar 2010) and the following options “-d gyrA_parC −c 0.95”. We did this both for the shotgun metagenomics data as well as for the amplicon sequences derived specifically from Enterobacteriaceae *gyrA* and *parC* genes. Prior to this sequence mapping raw reads were quality filtered using Trim Galore! (Babraham Bioinformatics 2012) with the settings “-e 0.1 −q 28 −O 1”. We then used the R script (R Core Team 2016) provided with the Mumame software to compare the matches to mutated and wildtype sequences in the database. The same database and method combination was used to quantify fluoroquinolone resistance mutations in sequence data from an Indian lake exposed to ciprofloxacin pollution (Bengtsson-Palme et al. 2014), as well as in an Indian river upstream and downstream of a wastewater treatment plant processing pharmaceutical waste (Kristiansson et al. 2011; Pal et al. 2016). These samples were preprocessed in the same way as in the Indian lake study (Bengtsson-Palme et al. 2014).

To quantify resistance mutations to tetracycline in the sequence data generated by (Lundström et al. 2016), we created a Mumame database for tetracycline resistance mutations in the 16S rRNA gene. We extracted the mutational information related to tetracycline from the CARD “snps.txt” file and then built the database using the following command: “mumame_build −i card-data/nucleotide_fasta_rRNA_gene_variant_model.fasta −m Tet_snps.txt −o Tet −n”. We then mapped all reads from the Lundström et al. (2016) data to the Mumame database using the options “-d Tet −c 0.95 −n”. Reads were quality filtered and statistical differences were assessed as above.

### Software evaluation

To assess the limitations of the method in terms of sequencing depth, the samples from the highest and lowest ciprofloxacin concentrations generated by Kraupner et al. (2018; 10 μg/L and 0 μg/L, respectively) were downsampled to 1, 5, 10, 20, 30, 40 and 50 million reads. Thereafter, the reads from the downsampled libraries were mapped to the fluoroquinolone resistance mutation database using Mumame as above. Statistical differences were assessed at all simulated sequencing depths and average effect sizes calculated for the significantly altered genes.

## Results

### Mumame can quantify point mutation frequencies in metagenomic data

As a proof-of-concept that our method to identify point mutations in metagenomic sequence data is functional, we used Mumame to quantify the mutations in amplicon data from the the *gyrA* and *parC* genes. These genes are targets of fluoroquinolone antibiotics, and often acquire resistance mutations attaining high levels of resistance. We quantified such mutations in an amplicon data set specifically targeting these two genes in *Escherichia coli*. This data set derives from an exposure study with increasing ciprofloxacin concentrations, and enrichments of mutations in the classical fluoroquinolone resistance determining positions S83 and D87 (*gyrA*) and S80 and E84 (*parC*) have previously been verified using other bioinformatic methods (Kraupner et al. 2018). This data set therefore serves as an ideal positive control for our novel method. We found that Mumame were able to identify the difference between the highest concentration (10 μg/L) and the lower ones reported in the original study (Figure 1). However, Mumame only reported an average frequency of mutations of around 11-12% for *gyrA* mutations (Figure 1A), while the original paper finds frequencies of 60-85% (S83) and 30-40% (D87). The A67 position was not quantified in the original paper. The reason for the discrepancies is unknown, but it is likely caused by a taxonomic filtration step that selects for *E. coli* reads used in the Kraupner et al. study, while Mumame does not perform prior filtering. The decision to exclude filtering was made in order to mimic a situation with true metagenomic data where several target species may co-exist. For *parC*, Mumame only quantified the S80 position (Figure 1C), because the E84 mutations were not included in the version of the CARD database used for this study. For position S80, Mumame identified around 35% mutated sequences at the highest concentration of ciprofloxacin, while the original study reported around 50%.

**Figure 1.**
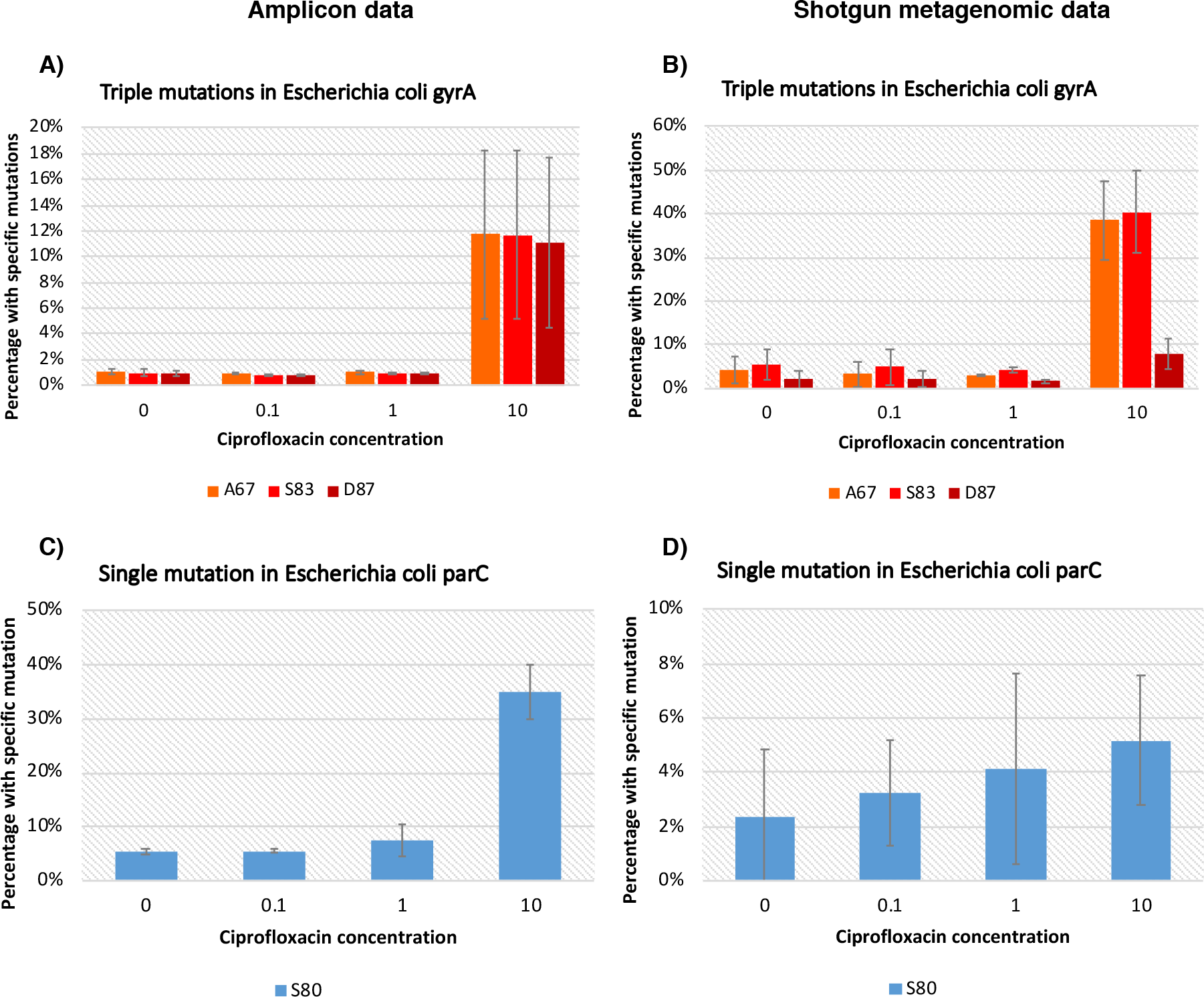
Total mutation frequencies quantified using Mumame for three known mutations conferring resistance to fluoroquinolone in the *E. coli gyrA* gene based on amplicon sequencing (A) and shotgun metagenomic data (B) from the same samples. Corresponding data for the S80 mutation in *parC* is shown in (C) for amplicon data and (D) for shotgun data.

We next evaluated the performance of Mumame on the real shotgun data that was also generated from the same samples as the amplicon libraries. Ideally, this analysis should generate virtually the same result as the amplicon analysis. Indeed, we found similar results for the A67 and S83 *gyrA* mutations (Figure 1B). For the D87 mutation, the frequencies were much lower than for the other two mutations, albeit still significantly larger than at the lower concentrations (p < 0.01).

For the *parC* gene, the shotgun metagenomic analysis was too noisy to generate a statistically significant result, which was highly surprising to us (Figure 1D). Taken together, these results indicate the high noise levels present for individual gene variants even in deeply sequenced shotgun metagenomes from controlled exposure studies.

### The limits to quantification

Noting the much more instable levels of mutations in the shotgun metagenomes, we next investigated the effects of sequencing depth on the ability of our method to detect significantly altered mutation frequencies. For this analysis, we used downsampled data from the shotgun metagenomic library of the ciprofloxacin exposure study (Figure 2). As expected, we found that the number of significantly altered mutation frequencies detected increased with larger sequencing depth (Figure 2A). In addition, the average effect size of the significant mutations became gradually lower with larger sequence depth, also in accordance with expectations (Figure 2B). Importantly, the average effect size of detectable mutation frequency differences seems to decrease linearly with sequencing depth. This means that we can calculate an expected detection limit for the method given the characteristics of the data and experimental setup. At 10 million reads, we expect that the proportion of reads with mutation must be 30-40% higher in the exposed sample in order for it to be detected as significant. The required effect decreases to, on average, 10% higher at 50 million reads (Figure 2B). These numbers are of course also dependent on other factors, such as the number of replicates per treatment, but nevertheless they can be used as ballpark numbers to aid the design of metagenomic studies or to interpret non-significant results derived from Mumame analyses.

**Figure 2.**
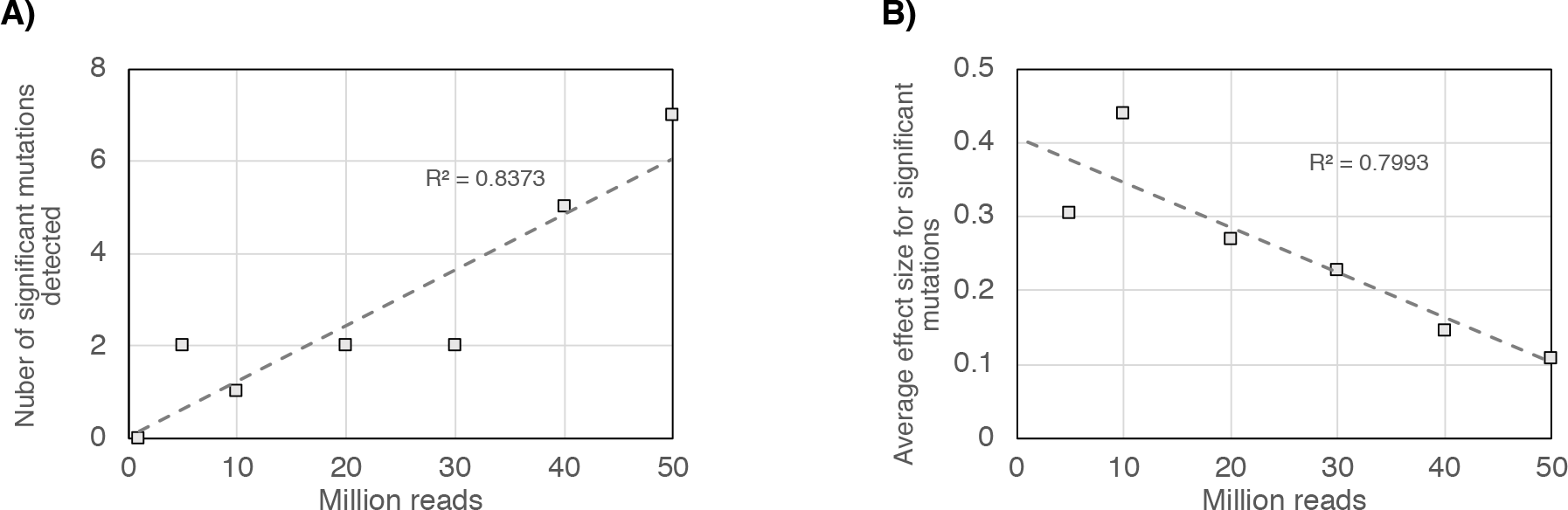
Relationship between the number of investigated reads and number of mutations with significantly altered frequencies (A) and the average effect size for those mutations (B); as assessed using Mumame on shotgun metagenomic data from a ciprofloxacin exposure experiment.

### Tetracycline-exposed Escherichia coli populations do not harbor higher abundances of resistance mutations

After performing the validation and limitation testing of the method, we next used Mumame to quantify resistance mutations in a similar controlled aquarium setup under exposure to the antibiotic tetracycline (Lundström et al. 2016). In this study, no amplicon sequencing of the target gene for tetracycline – the 23S rRNA – was performed, and thus there was no *a priori* true result that we could compare to. While Mumame was able to successfully detect tetracycline resistance mutations in the data, we somewhat surprisingly found no enrichment of tetracycline resistance mutations in this data (Figure 3). Notably, this result was obtained despite a very high sequencing depth (on average 181,595,072 paired-end sequences per library). Obtaining a negative result at this sequencing depth suggests that there actually is no enrichment of known *E. coli* resistance mutations in the samples.

**Figure 3.**
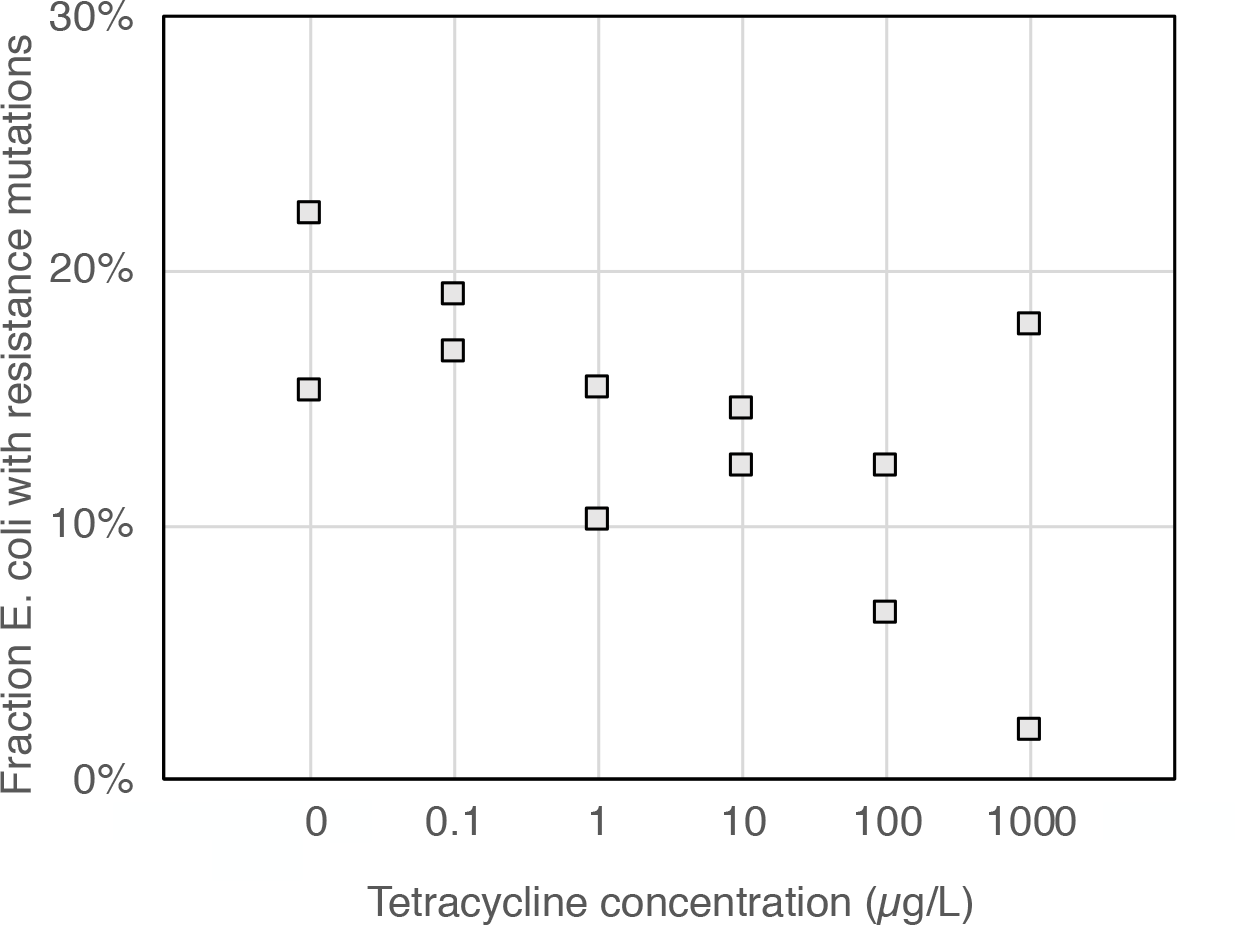
Frequencies of *E. coli* tetracycline resistance mutations at exposure to different concentrations of tetracycline, based on shotgun metagenomic data.

### Fluoroquinolone resistance mutations in ciprofloxacin-polluted sediments

As a final investigation of the performance of the method, we also let Mumame quantify the fluoroquinolone resistance mutations in river and lake sediments polluted by antibiotic manufacturing waste, primarily ciprofloxacin (Kristiansson et al. 2011; Bengtsson-Palme et al. 2014; Pal et al. 2016). These libraries are fairly old and were not as deeply sequenced as the other data sets we investigated. While the experimental setup of these studies in terms of number of samples does not allow for proper statistical testing, we did find an enrichment of the fluoroquinolone resistance mutation frequencies downstream of the pollution source, at least for the *E. coli gyrA* and *parC* genes (Figure 4). We also detected a few such mutations in other species, but the counts of those were low and the results largely non-informative due to the small number of detections per mutation (Figure 5).

**Figure 4.**
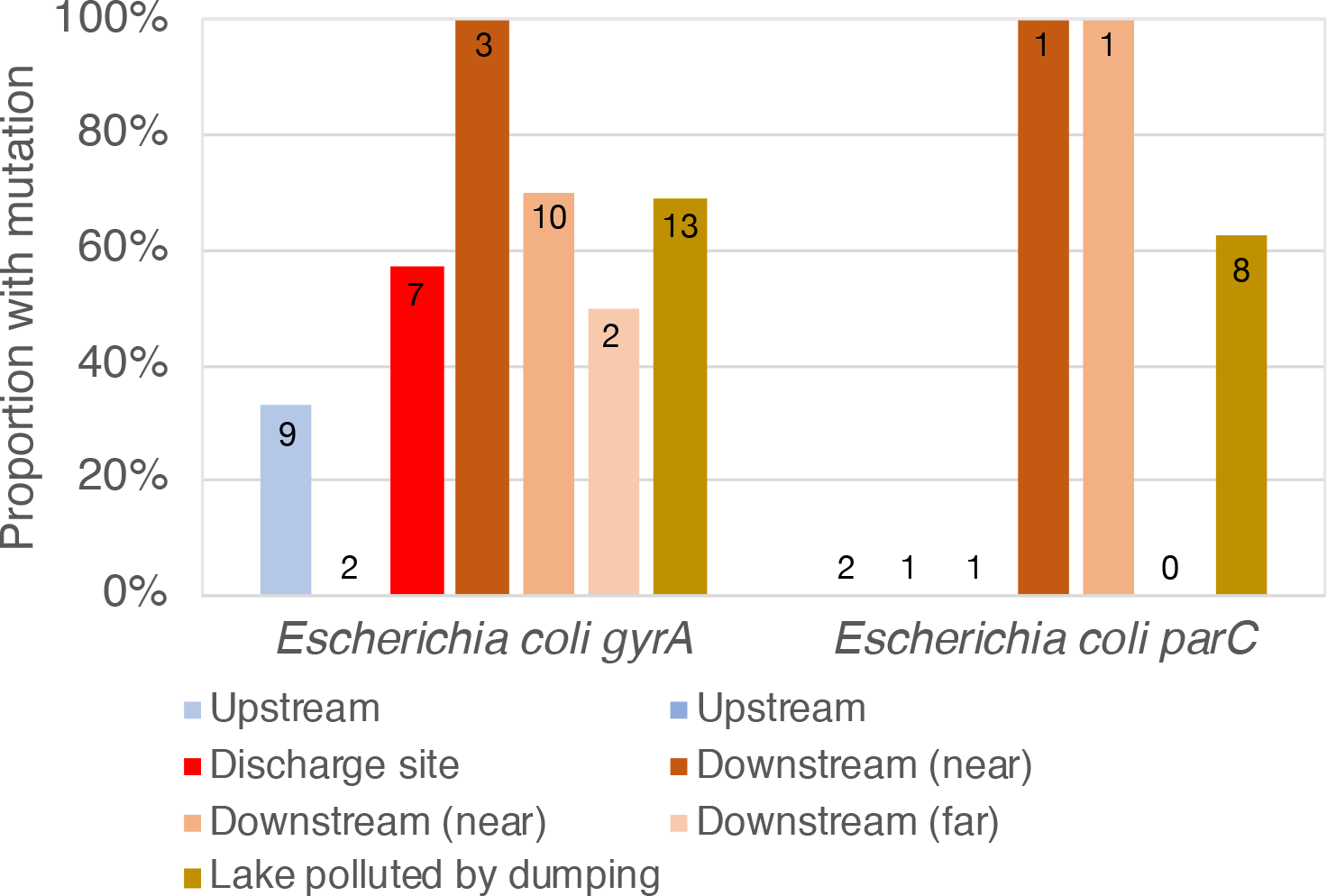
Relative frequency of *gyrA* and *parC* sequences with resistance mutations in samples taken downstream, at or upstream of the pharmaceutical production wastewater treatment plant, as well as in a lake polluted by dumping of pharmaceutical production waste. The numbers at the top of the bars shows the total number of sequences (wildtype or mutated) identified in each sample.

**Figure 5.**
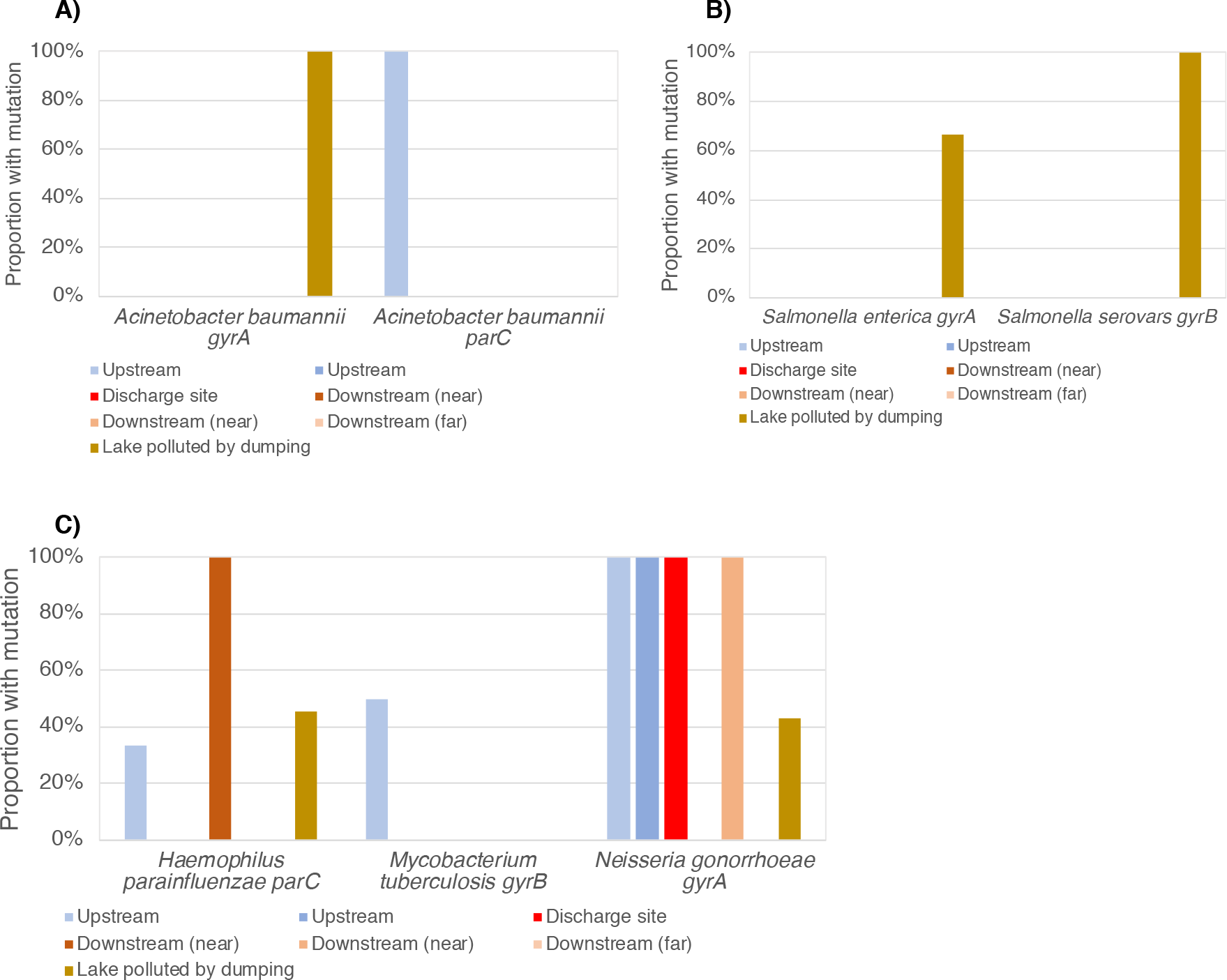
Relative frequency of sequences with resistance mutations in samples taken downstream, at or upstream of the pharmaceutical production wastewater treatment plant, as well as in a lake polluted by dumping of pharmaceutical production waste, for *Acinetobacter baumanii* (A), *Salmonella* species (B) and *Haemophilus parainfluenzae*, *Mycobacterium tubercolosis* and *Neisseria gonorrhoeae* (C).

## Discussion

Metagenomics often becomes restricted to investigate gross compositional changes to the taxonomy and function of microbial communities. Unfortunately, this obscures important variation between individual sequence variants that may have large outcomes on phenotypes (Österlund et al. 2017; Bengtsson-Palme 2018). One example of such point mutations inducing strong phenotypic changes is resistance mutations in the target genes of antibiotics (Kraupner et al. 2018). However, including mutated sequence variants in the antibiotic resistance gene databases is complicated, and can lead to gross misinterpretations of the data (see for example (Ma et al. 2014). Still, understanding relevant variation between sequences and linking that to phenotypes is somewhat of a holy Grail of metagenomics. This study has made clear that we are not yet at that point in terms of bioinformatic methods and the sequencing depth required to draw firm conclusions. That said, we show in this work that identifying significant and relevant differences in resistance mutation frequencies between sample groups from shotgun metagenomic data is possible, given a sufficiently large sequence depth. However, the quantitative estimates still seem to be highly variable, even at very large sequencing depths.

The results of the Mumame evaluation also provides a few other important clues on potential pitfalls with inferring mutation frequencies from shotgun metagenomic data. An important such aspect is the disparity between mutation frequencies described by amplicon sequencing and shotgun data. Particularly, the ability to relatively consistently identify the A67 and S83 mutations in *parC*, while the D87 mutation is seemingly less frequent in the shotgun data is somewhat troubling if the goal is to identify the actual abundances of such mutations. At the same time, the statistical significance of those differences could still be identified. For the A67 and S83 mutations, only 5 million reads were required for a significant effect to be detected, while for the D87 mutations a sequencing depth of 50 million reads was required. This is not necessarily a shortcoming of the Mumame software, but may just as well be due to the much noisier nature of counts from metagenomic sequence data compared to the large number of reads corresponding to the same genes deriving from amplicon data (Jonsson et al. 2016a).

Another important potential problem highlighted by our evaluation is the need to produce very large sequence data sets to be able to identify and quantify mutations (and wildtype) sequences with any certainty. As a rule of thumb, the targeted regions represent less than 0.004% of the bacterial genome, and each bacterial strain may correspond to only a fraction of a percent of the reads in the shotgun sequence data (depending on its abundance). This means that to identify a single read from a resistance region in the data, one would – on average – need to sequence more than five million reads. To get a reasonably confident measure of reads stemming from wildtype versus strains with mutations, approximately 10 reads from each group would be needed per sample (or, say, 20 reads in total). That would, as a rough estimate, correspond to a hundred million reads per sample. This is, unfortunately, way more sequences than what is typically generated per sample by shotgun metagenomic sequencing projects. In this study, only the samples from the tetracycline exposure study corresponded to such a high sequencing depth. Naturally, these numbers would depend on the proportions of the targeted microorganisms as well as their genome sizes, but ultimately this still presents the largest limitation to mutation studies based on metagenomic sequence data. Potentially, this problem could be partially alleviated by analyzing sufficiently large cohorts and perform the statistical analysis for general trends, but even large cohorts would be insufficient for mutations rare enough to pass below the detection limit.

In terms of interpreting the results from the exposure experiments, it is interesting to note the overall clear increase of fluoroquinolone resistance mutations at the highest ciprofloxacin concentration, which nearly perfectly correspond to increases in mobile *qnr* fluoroquinolone genes in the same samples (Kraupner et al. 2018). This is contrasted by the trend seen in the tetracycline exposure experiments, where tetracycline resistance genes – specifically efflux pumps – were enriched at higher tetracycline concentrations (Lundström et al. 2016), while tetracycline resistance mutation abundances were not significantly altered. This non-significant result was obtained despite the exceptionally high sequencing depth of those samples.

While we did not have data from a proper experimental setup to address differences between sediments exposed to different degrees of fluoroquinolone pollution, the quantification of resistance mutations seems to provide an important piece of information to explain the results of previous studies of resistance gene abundances in these river samples (Kristiansson et al. 2011). In the original paper, the abundance of mobile fluoroquinolone resistance genes (*qnr* genes) were shown to be enriched in the low-level polluted upstream samples, compared to the highly polluted downstream samples. Importantly, the *qnr* genes only provide resistance to relatively low levels of fluoroquinolones (Hooper and Jacoby 2015), and the authors of hypothesize that chromosomal mutations of the target genes are probably necessary to survive the selection pressure from antibiotics downstream of the pollution source. In this work, we show that this assumption is likely correct. Only a limited number of reads were mapping to these resistance regions and the number of samples unfortunately prevents us from properly assessing a statistical difference between the upstream and downstream samples. Still, the proportion of resistance mutations seems to be systematically higher in the samples downstream of the pollution source, at least for *E. coli*. This indicates that the method we present here can provide important additional information to metagenomic studies of resistance patterns in different environment types, given that a sufficient sequencing depth is achieved.

We have here shown the utility of the Mumame tool for finding resistance mutations in shotgun metagenomic data. In this paper, we have used the CARD database (Jia et al. 2016) as the information source for resistance mutation, but the tool is flexible to use any source of such data. It is also not in any means restricted to the mutations investigated in this paper but is fundamentally agnostic to the input data. It can also be used in open screening for mutations in any gene present in the database in parallel, and can handle different mutations in both RNA and protein coding genes. The tool is flexible and fast and can therefore be implemented as a part nearly any screening pipeline for antibiotic resistance data in metagenomic data sets.

## Conclusion

This paper presents a software tool called Mumame to analyze shotgun metagenomic data for point mutations, such as those conferring antibiotic resistance to bacteria. Mumame distinguish between wildtype and mutated gene variants in metagenomic data and quantify them, given a sufficient sequencing depth. We also provide a statistical framework for handling the generated count data and account for factors such as differences in sequencing depth. Importantly, our study also reveals the importance of a high sequencing depth – preferably more than 50 million sequenced reads per sample – in order to get reasonably accurate estimates of mutation frequencies, particularly for rare genes or species. The Mumame software package is freely available from http://microbiology.se/software/mumame. We expect Mumame to be a useful addition to metagenomic studies of e.g. antibiotic resistance, and to increase the detail by which metagenomes can be screened for phenotypically important differences.

## Acknowledgements

This work was funded by the Swedish Research Council for Environment, Agricultural Sciences and Spatial Planning (FORMAS; grant 2016-00768).

## Author contributions

JBP conceived of the study. SM and JBP collected and analyzed the data. JBP designed and wrote the software package. VJ provided statistical guidance for the R implementation. JBP wrote the draft manuscript. All authors interpreted the data and contributed to the writing of the paper.

